# Status of Florida’s pillar coral population: in situ declines and ex situ successes

**DOI:** 10.1101/2025.06.19.660574

**Authors:** Karen L. Neely, Rachel M. Morgan, Keri O’Neil, Nikkie Cox, David Gilliam, Michelle Mair, Brian Walker, Amanda Zummo, Cailin Harrell, Ananda Ellis

**Affiliations:** National Coral Reef Institute, Nova Southeastern University, Dania Beach FL, USA; The Florida Aquarium Coral Conservation and Research Center, Apollo Beach FL, USA; Coral Restoration Foundation, Tavernier FL, USA; Florida Fish & Wildlife Conservation Commission, Fish and Wildlife Research Institute, Marathon FL, USA

**Author notes:** **Correspondence** Karen L. Neely.

**Keywords:** SCTLD, bleaching, population decline, pillar coral, propagation, spawning

## Abstract

The population of the pillar coral, *Dendrogyra cylindrus*, in Florida was decimated from 2013-2020, primarily by the emergence of stony coral tissue loss disease (SCTLD). Monitoring of survivors from 2021 – early 2025 showed that the population underwent an additional 96% decline in live tissue, 78% loss in living colonies, and 57% loss of genotypes. SCTLD continued to be the primary cause of these losses. Though some surviving tissue isolates exhibited small amounts of growth, the population remains extremely small, with only an estimated 9.6 square meters of tissue remaining on 23 colonies (15 genotypes). Additionally, colonies are far too dispersed to successfully fertilize spawned gametes. The further declines in the population since 2020 highlight the instability of the remnant population, as well as the value of the pillar coral rescue program and ongoing propagation efforts. As of February 2025, eight different in situ and ex situ facilities were caring for rescued *D. cylindrus*. Experimental fragmentation at one in situ nursery identified variable, but continually increasing, growth rates across multiple fragmentation events. Sexual propagation efforts at an ex situ nursery documented 105 different rescue fragments spawning across five years. The smallest fragment was 9 × 7 × 9 cm, establishing a potential “minimum colony size” for reproductive capacity for this species. From these spawning events, 82 juveniles were being raised ex situ in early 2025. Two of these sexually propagated juveniles spawned six years after settlement, thus establishing a potential minimum age for reproduction.

## Introduction

The pillar coral *Dendrogyra cylindrus* is a historically uncommon but conspicuous coral in the Caribbean region. It is listed as critically endangered on the IUCN Red List (Aronson et al. 2008), and endangered under the U.S. Endangered Species Act. The Florida population of the species was extensively monitored from 2013-2020 (Jones et al. 2021; Neely et al. 2021a). During that time, the population suffered extensive decline, with minor losses from bleaching and white plague, and unprecedented losses from stony coral tissue loss disease (SCTLD). Losses to the population through the end of 2020 included 94% loss of tissue, 93% of colonies, and 86% of genotypes (Neely et al. 2021a).

This catastrophic decline resulted in the functional extinction of the species in Florida, with only a handful of survivors scattered across the reef tract and essentially no chance for successful reproduction in the wild. From 2021-2025, we continued to monitor the remaining colonies for continued declines or potential recovery, and also updated the Florida *D. cylindrus* database when previously unknown colonies (live or dead) were found.

The rapid decline of *D. cylindrus* in Florida also prompted a rescue program in which over 550 fragments of a presumed 128 wild genotypes were collected between 2015 and 2019 and brought to in situ and ex situ facilities for care (Neely et al. 2021b). These corals have grown substantially and reproduced successfully (O’Neil et al. 2021), and we here provide updates on both sexual and asexual production of rescued individuals.

## Methods

### Newly discovered corals

Reports of *D. cylindrus* colonies submitted to the authors were cross-referenced with existing geographic coordinates and photos to determine whether their presence was already documented. If not, coral size (straight line length, width, and height), percent coverage of live tissue, and depth were added to the database. For any sites with multiple colonies, distances and bearings were taken between corals for differentiation and repeatable identification of individuals.

### Monitoring of survivors

Surviving *D. cylindrus* were monitored at least annually. Colonies in Dry Tortugas National Park were monitored tri-annually, those in southeast Florida were monitored monthly or tri-annually, and those in Florida Keys National Marine Sanctuary were monitored opportunistically at least once a year but sometimes more frequently. During each monitoring event, corals were assessed for percent live cover and percent recent mortality, with the cause(s) of any recent mortality recorded.

### Calculating population change

The amount of live *D. cylindrus* tissue on Florida reefs was calculated by multiplying the estimated surface area of the entire colony (assumed and calculated as a half ovoid using length (L), width (W), and height (H) measurements) by the proportion of living tissue remaining on each.

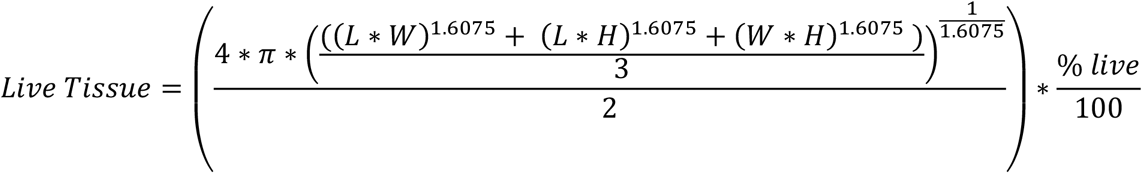

The amount of tissue remaining at the beginning of 2025 was compared to the amount remaining at the end of 2020, as well as the amount of tissue present during the 2013-14 baseline surveys. Any living *D. cylindrus* colonies found after the baseline surveys were assumed to have remained unchanged since 2013. Given that these corals very likely underwent tissue losses since 2013-14 (as observed via long-term colony monitoring), our estimates are conservative, with actual tissue losses probably much higher.

The proportion of colonies remaining in 2025 was calculated by dividing the number of colonies alive in early 2025 by the number known to be alive at two previous time points: the end of 2020, and the 2013-14 baseline. Any dead colonies first found after 2020 were assumed to have been dead since the baseline, again making the reported losses conservative.

The proportion of genotypes remaining in 2025 was calculated by dividing the number alive in early 2025 by the number known to be alive at the end of 2020 and the number known to be alive at the baseline surveys. As described in Neely et al. (2021a), a genotype was defined as a colony or group of colonies specifically genotyped in Chan et al. (2019) or, when colonies were not genotyped, any colony which was at least 70 m from any other colony.

We also replicated the methods from Neely et al. (2021a) when estimating the distances between *D. cylindrus* genotypes to highlight the unlikelihood of successful sexual reproduction via unassisted spawning. We used ArcGISPro to calculate the nearest surviving genotype for each genotype.

### Rescue population breeding and settler growth: land-based nursery

Induced spawning and propagation efforts for ex situ *D. cylindrus* have been ongoing at The Florida Aquarium since 2019. We assessed the total number of distinct fragments observed spawning across all years, and identified the smallest spawning individuals.

To assess growth, we measured the diameter of each surviving settler from each year’s spawning cohort. Measurements were taken in February 2025 for surviving settlers from 2018, 2021, 2022, and 2024 spawning events.

### Rescue population fragmentation and growth: field-based nursery

*Dendrogyra cylindrus* fragments historically collected from wild colonies were housed on “boulder tree” platforms – PVC trays with fixed mounts – suspended in the water column at the Coral Restoration Foundation nursery in the upper Florida Keys. Fragments were asexually propagated with the goal of creating additional fragments for eventual outplanting as well as increasing growth rates. Asexual propagation was conducted across four time periods. In January 2021, fragments were created in situ using hand cutters and loppers or, for larger *D. cylindrus* pieces, temporarily moved ex situ and cut with a wet tile saw. In January 2022, in situ fragments were cut the same way while ex situ fragments were cut with a wet band saw. In two additional fragmentation events – November/December 2022 and February 2023 – all fragments were cut in situ. All fragments were epoxied to PVC cards and returned to the boulder trees.

The maximum diameter of each fragment from the January 2021 – January 2022 propagation events was measured regularly using calipers. Fragments produced from the November 2022 – February 2023 events were measured regularly using top-down photographs that were assessed for total surface area using ImageJ and then recalculated for diameter using the assumption of a circular area. The change in maximum diameter of each fragment between the asexual propagation event and the end of monitoring (147 – 539 days) was calculated. Any corals for which partial or full mortality was observed, or for which the end maximum diameter was smaller than the initial, were discarded from the dataset as non-representative of normal growth rates. A multiple regression was used to assess correlations between date of fragmentation (with dummy values for each of the four fragmentation events) and maximum diameter of the newly cut fragments to the growth rates.

## Results

### In situ colonies

The total number of known *D. cylindrus* colonies, live or dead, on Florida’s Coral Reef as of February 2025 was 876 (Figure 1). A total of 811 of these (93%) were alive upon discovery. Of the 876 colonies, 61 of them, representing a presumed 13 genotypes, were discovered since the publication of Neely et al. (2021a). Of these new discoveries, 46 colonies (3 new genotypes) were found in Dry Tortugas National Park as part of unprecedented coral survey efforts ahead of the arrival of SCTLD to the area. The Dry Tortugas colonies were all alive at first sighting, but of the colonies found elsewhere on Florida’s Coral Reef from 2021 – 2025, only 5 of the 15 colonies had live tissue. Two of these, both in Biscayne National Park, had all remaining tissue collected for ex situ rescue in 2023.

**Figure 1.**
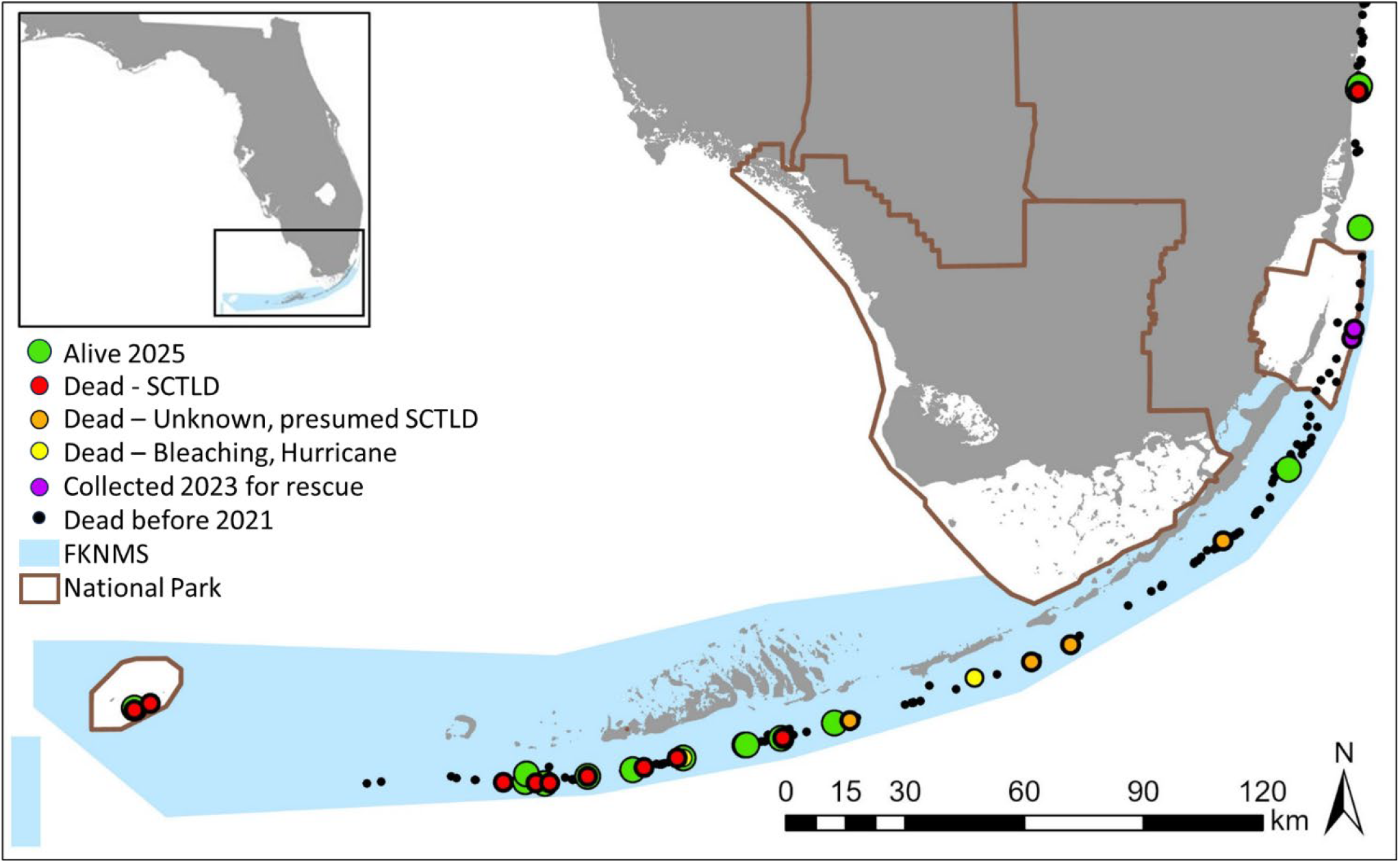
Distribution of *Dendrogyra cylindrus* genotypes on the Florida Reef Tract. Most colonies were dead before 2021 (black dots). Of the genotypes still alive at the end of 2020, by 2025 some retained live tissue (green dots), some died of stony coral tissue loss disease (red dots), some died of presumed SCTLD (orange dots), two died from other causes (one from bleaching, one from a hurricane; yellow dots), and two had their last remaining tissue collected for ex situ holding (purple dots). All corals lie within (from southwest to northeast) Dry Tortugas National Park, Florida Keys National Marine Sanctuary, Biscayne National Park, or the southeast Florida region.

Monitoring of survivors from 2021-2025 documented continued declines in coral tissue, living colonies, and surviving genotypes (Figure 2A). From 2021 – 2025, the amount of living *D. cylindrus* tissue on Florida’s Coral Reef declined an additional 96%, for a total loss of at least 99.7% since 2013 baseline surveys. The number of surviving colonies also continued to decline, with a 78% loss of colonies between 2021 and 2025, for a total loss of 97% since baseline surveys. Similarly, the number of surviving genotypes declined an additional 55% from 2021-2025 for a total loss of 92% since baseline surveys.

**Figure 2.**
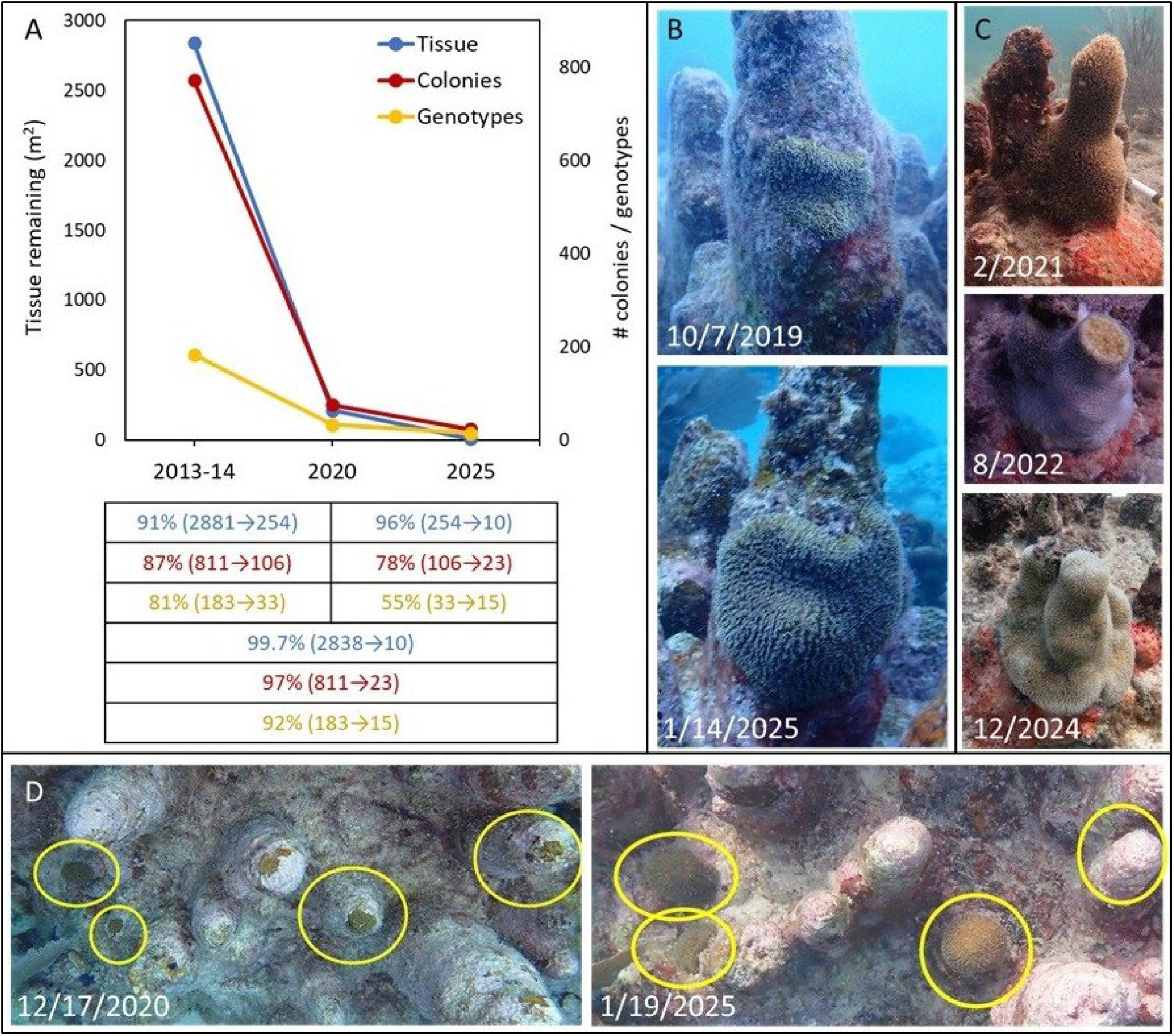
A) Decline of the Florida *Dendrogyra cylindrus* population as measured by square meters of live tissue (blue), number of living colonies (red), and number of surviving genotypes (yellow). Values in the table show the changes in each metric from the 2013-14 baseline surveys to the end of 2020, from the beginning of 2021 to early 2025, and from the baseline surveys to early 2025. Values from baseline to 2020 differ slightly from Neely et al 2021 because newly found colonies were included in these updated numbers. B) Growth of the last remaining tissue isolate of a Lower Keys genotype. C) Changes to a tissue isolate in the southeast Florida region, showing the loss of the pillar tip to an unknown cause in 2022 followed by growth of the isolate. D) Four remaining isolates (yellow circles) of a Lower Keys genotype, showing growth in three of them, but death of the fourth.

By the beginning of 2025, we estimated only 9.6 m^2^ of living *D. cylindrus* tissue, located on 23 colonies representing 15 genotypes, on Florida’s reefs. Of this tissue, 26% existed within Dry Tortugas National Park (8 colonies, 1 genotype), 24% within the Florida Keys National Marine Sanctuary (11 colonies, 11 genotypes), and 49% on southeast Florida reefs north of Biscayne National Park (4 colonies, 3 genotypes). All surviving colonies underwent substantial tissue loss over the years. On the survivors, the percentage of tissue remaining by 2025 ranged from 0.1 – 25%. With only three exceptions (10%, 15%, and 25% - all in the southeast Florida region), all survivors had only 5% or less of their tissue remaining.

For genotypes remaining in 2025, the median distance between survivors was 8.0 kilometers. Only two genotypes resided within less than 1 km (2 at 402 meters) of a conspecific. The most remote genotype was the sole surviving genotype in Dry Tortugas National Park, which was 91.4 km from its nearest neighbor.

Losses to the *D. cylindrus* population from 2021-2025 continued to be predominantly caused by stony coral tissue loss disease (SCTLD). Of the 18 genotypes lost during that time, 12 were observed perishing completely from SCTLD lesions, and an additional 4 were presumed as such based on appearance and absence of other stressors. One genotype was lost to a hurricane which knocked off the last remaining piece of tissue. Another genotype, which had 12 individual colonies with varying degrees of live tissue in early summer of 2023, experienced total losses in live tissue by November 2023 from bleaching-related mortality. All other *D. cylindrus* genotypes in Florida bleached completely during this hyperthermal event but had no resulting tissue loss.

The surviving corals generally have only one or a few small isolates of live tissue remaining. While most colonies have continued to lose tissue since 2020, we know of at least three colonies where isolates have shown moderate growth (Figure 2B-D).

### Rescue population breeding and settler growth: land-based nursery

In total, 105 *D. cylindrus* fragments rescued from wild colonies and held at The Florida Aquarium Coral Conservation and Research Center (CCRC) spawned over five years of propagation efforts. The smallest ex situ fragment observed spawning was 9 × 7 × 9 cm (L x W x H). In 2024, spawning efforts focused on fragments greater than or equal to that, and 75% of those successfully spawned.

In February 2025, sexual recruits that settled six months prior (2024 cohort) measured between 1.5 – 4 mm in maximum diameter (Figure 3). Recruits from 42 and 54 months earlier (2020 and 2021 cohorts) ranged from 6-8 cm. And those spawned 78 months earlier (2018 cohort; spawned at Keys Marine Lab from wild-collected individuals and transported to CCRC) ranged from 12-15 cm.

**Figure 3.**
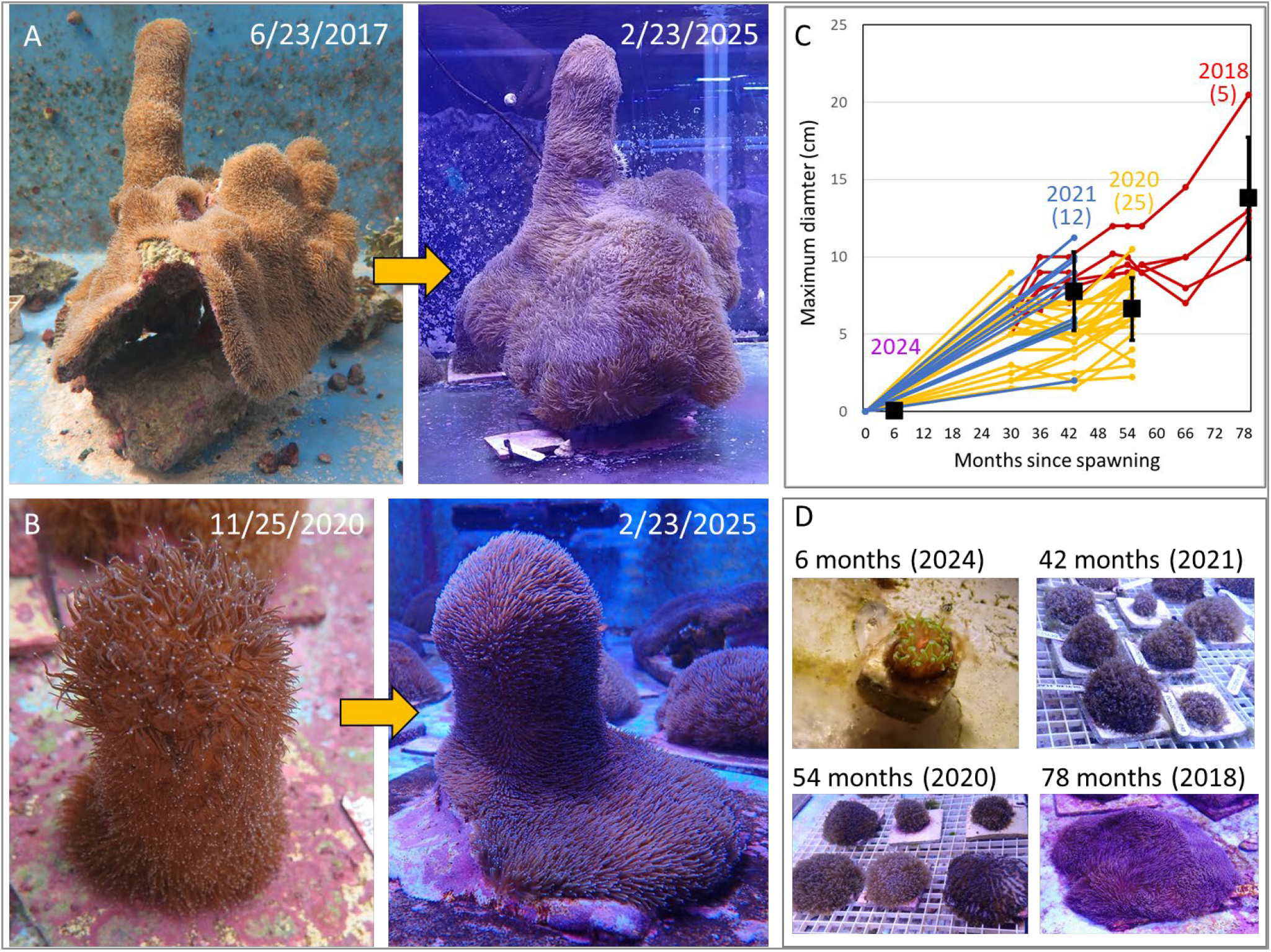
(A,B) Examples of multi-year growth of two *Dendrogyra cylindrus* fragments collected from the wild and held in ex situ raceways. (C) Size of sexually produced juveniles from four cohorts (2018, 2020, 2021, and 2024). Individual lines represent growth measurements of specific juveniles, while black boxes represent the mean maximum diameters (± SD) of corals from each cohort as measured in February 2025. (D) Example images taken in February 2025 of sexually propagated juveniles from the four different year cohorts.

Assuming a linear growth rate, the maximum diameter of sexually spawned individuals increased at 1.8 mm/month from settlement to 6.5 years of age. Three settlers from the 2018 sexually produced cohort spawned as males in 2024 (sizes at spawning: 9.5 × 8 × 8, 10 × 10 × 5.5cm and 14.5 × 13 × 5cm). Rescue project fragments collected in situ and held at the CCRC also exhibited notable growth (Figure 3).

### Rescue population fragmentation and growth: field-based nursery

The maximum diameter growth rate of *D. cylindrus* fragments at Coral Restoration Foundation’s nursery varied across fragments from 0 – 3.58 mm/month (Figure 4), with an average of 0.83 (± 0.68) mm / month. A multiple linear regression (R^2^ = 0.31) found that the maximum diameter of a newly-cut fragment was not a significant factor for subsequent growth rates (p = 0.16). Growth rates did, however, increase with each subsequent fragmentation date (January 2021: 0.44 ± 0.33. January 2022: 0.65 ± 0.43. November/December 2022: 1.04 ± 0.68. February 2023: 1.45 ± 0.88 mm/month), and the multiple regression identified fragmentation date as a significant factor in determining growth rate (January 2021: p < 0.001. January 2022: p < 0.001. November/December 2022: p = 0.02).

**Figure 4.**
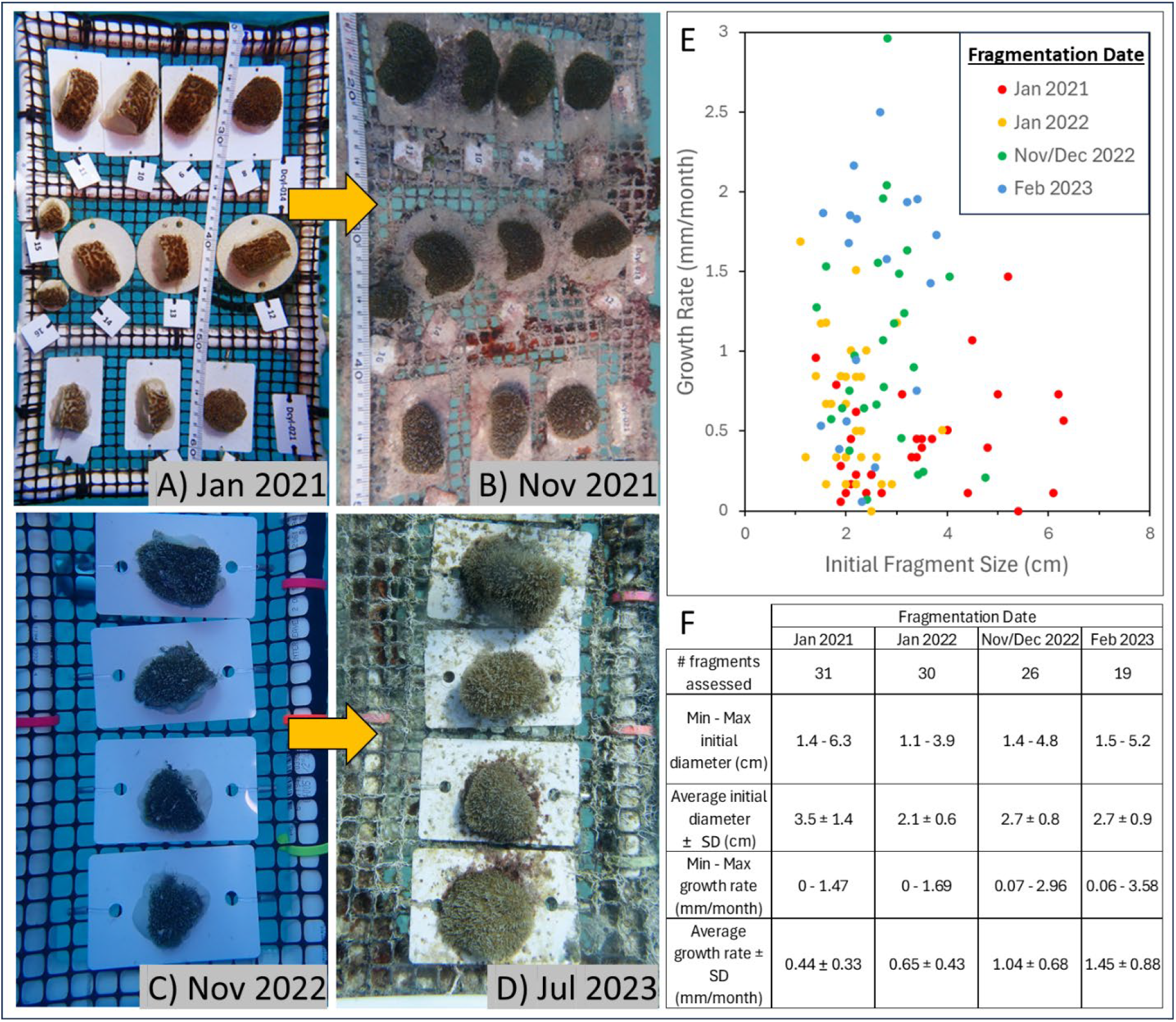
(A, C) Examples of fragment growth between the initial fragmentation dates and (B,D) subsequent monitoring dates. (E, F) Growth rates were not equal across fragmentation events, with later ones having faster growth rates than earlier ones.

## Discussion

The discovery of a small number of previously unknown *D. cylindrus* genotypes from 2020-2024 indicates that there may still be other survivors. However, the slow rate of discovery, despite extensive surveys conducted by various monitoring and coral restoration programs in Florida each year, combined with the small number of colonies found alive, indicates that the vast majority of the corals of this species in Florida have already been found. It is highly unlikely that there is an unknown remnant population.

The known remaining individuals are incapable of rebuilding the population. As of February 2025, only nine of the surviving 23 individuals had tissue isolates meeting the minimum observed spawning size. Additionally, the median distance between surviving genotypes increased from the already large 1.0 km in 2021 to over 8.0 km in 2025. The two closest remaining genotypes (403 m apart) both contain only a few square centimeters of remaining tissue.

SCTLD remains the biggest threat to this species in Florida, where the disease is largely considered endemic. Of the losses to genotypes since 2020, 89% died from SCTLD. All surviving colonies in Dry Tortugas and two of those within FKNMS have been kept alive through amoxicillin treatments on SCTLD lesions. These treatments are no longer authorized within FKNMS, so further losses are expected. Water temperatures and cumulative thermal stress in 2023 far exceeded those of any prior year (reviewed in Neely et al. 2024). Despite all remaining *D. cylindrus* colonies fully bleaching during that time, only a single genotype was lost, indicating that the remaining colonies are capable of handling temperature extremes and are unlikely to face mortality from this particular threat unless new thermal stress records are set.

Despite the dire situation of the wild population, husbandry, fragmentation, and propagation efforts have demonstrated that ex situ corals can grow and reproduce. Fragmentation of rescued individuals identified highly variable growth rates, with increases in diameter during the 2023 fragmentation event averaging over three times greater than those from the 2021 one. This variability may have resulted from changing environmental factors through time within the nursery, minor changes in handling during fragmentation, improved resilience of pre-fragmented colonies with longer pre-fragmentation nursery time, or other unknown factors. Further experimentation into optimizing growth rates based on these factors are needed to improve the efficiency of propagation efforts. *D. cylindrus* fragments at Coral Restoration Foundation were generally cut to approximately 2.5 cm in diameter. With an average growth rate of 0.83 mm/ month, we estimate that fragments would need to be held within the nursery for an average of 6.5 years to reach the minimum spawning size (9 cm) observed by The Florida Aquarium. Nurseries may need to plan for extended durations for fragments of this species to reach maturity.

Ex situ sexual propagation has resulted in successful larval production and settlement. Juvenile rearing requires extensive husbandry, yet successful spawning at age six by sexually produced juveniles indicates potential for multi-generational ex situ production. The genetic material for this population, including next-generation individuals, remains extant, and it is hoped that one day these genotypes and their offspring may be returned to Florida’s Coral Reef.

## Acknowledgments

We are grateful to the past and present divers of the following Nova Southeastern University labs: Disease Intervention, Coral Reef Restoration, Assessment & Monitoring (CRRAM), and GIS/Spatial Ecology. In particular, we thank Michelle Dobler, Kevin Macaulay, Arelys Chaparro, Kathryn Toth, and Reagan Sharkey. We also thank Amelia Moura, Phanor Montoya-Maya, as well as staff and interns involved in the pilot study at Coral Restoration Foundation.

## Data Availability Statement

The raw data supporting the conclusions of this article will be made available by the authors, without undue reservation.

